# Hypoxia increases neural proliferation, alters vascular structure, and reprograms the transcriptome and proteome of the speckled sanddab brain

**DOI:** 10.64898/2026.06.03.729957

**Authors:** Zurine De Miguel, Paul Stephens, Arie Dash, Griffin Bohman, Alina Diez, Alexandra Stella, Cheryl A. Logan, Scott L. Hamilton

## Abstract

Hypoxia (low oxygen availability) is a common environmental stressor in estuarine ecosystems that negatively affects fish survival as well as physiological and behavioral responses. However, the effects of hypoxia on the brain remains poorly understood, particularly in non-model species. Here, we investigated how prolonged hypoxia influences neural, vascular, and molecular responses in the brain of the speckled sanddab (*Citharichthys stigmaeus*), an ecologically relevant estuarine flatfish. Fish were exposed to normoxic or hypoxic conditions for seven days, and responses were assessed using histological analyses of neural proliferation and vascular structure, alongside transcriptomic and proteomic profiling. Hypoxia increased neural cell proliferation and progenitor activation in the hypothalamic nucleus recessus lateralis (NRL) and optic tectum, while reducing survival of newly generated cells. At the tissue level, hypoxia induced region-specific vascular remodeling, characterized by increased vessel area and vessel number without evidence of widespread endothelial proliferation. At the molecular level, transcriptomic and proteomic analyses revealed consistent enrichment of biological processes related to stress responses, development, metabolism, and cellular homeostasis, despite limited overlap between individual genes and proteins. Gene- and protein-level analyses further indicated activation of hypoxia-responsive pathways, including HIF signaling and oxidative stress protection, alongside selective metabolic reprogramming. Together, these findings demonstrate that hypoxia induces multi-level changes in the brain, linking neural plasticity, vascular remodeling, and molecular responses. This integrated response likely supports brain function under reduced oxygen availability in dynamic estuarine environments and highlights the role of the brain in regulating responses to environmental stress.

## Introduction

Hypoxia is a widespread environmental stressor in marine and estuarine ecosystems, driven by climate change, eutrophication, and tidal mixing processes (1, 2). Events of prolonged hypoxia (defined as < 2 mg O₂/L (3–5)) have become more frequent and severe in coastal habitats, including the Elkhorn Slough estuary in central California, where some fish populations experience chronically low oxygen conditions (6, 7). These environmental changes have been linked to reduced growth, survival, and reproduction in estuarine fishes, with cascading effects on population dynamics and ecosystem structure (6, 8–11). Estuaries are often important nursery habitats for many fish species (12, 13) and increasing hypoxia may threaten that nursery function, with ensuing impacts on offshore fisheries.

In fishes, hypoxia tolerance arises from coordinated molecular, physiological, and behavioral adjustments that conserve energy and maintain oxygen delivery (11, 14–17). At the molecular level, one of the best-characterized responses is the hypoxia-inducible factor (HIF) signaling pathway, which regulates gene expression programs involved in angiogenesis, metabolism, and cellular stress responses (18–22). In fish, multiple HIF-α isoforms (e.g., HIF-1α, HIF-2α, HIF-3α) show distinct expression patterns and may contribute to species-specific hypoxia tolerance mechanisms (18, 23). While important progress has been made in characterizing hypoxia responses in tissues such as gill, spleen, liver, muscle, and blood (24–31), comparatively fewer studies have examined hypoxia-responsive molecular programs in the fish brain (32–36).

The brain plays a central role in coordinating the environmental stress response, integrating sensory input and regulating physiological and behavioral adaptations to environmental challenges (37–39). In fishes, oxygen-sensitive neurons in the brainstem and hypothalamus detect changes in dissolved oxygen and initiate ventilatory and cardiovascular responses to cope with the stressor (40, 41). Neuroendocrine pathways such as the hypothalamus–pituitary–interrenal (HPI) axis regulate cortisol release, driving metabolic and behavioral responses to hypoxia (42), while modulation of the hypothalamus–pituitary–gonad (HPG) axis links hypoxia to reproductive regulation (43–45). However, at the same time, the fish brain is among the most vulnerable tissues to hypoxia, due to its high metabolic oxygen demand and reliance on aerobic ATP production (46). Reduced oxygen availability lowers mitochondrial ATP production and ion homeostasis (46), and alters neuronal excitability, synaptic plasticity, and cognitive function (47, 48). These processes are frequently accompanied by increased oxidative stress in brain tissue (49). Under hypoxia, cerebral metabolism shifts away from oxidative phosphorylation toward anaerobic ATP production (46) alongside broader metabolic reprogramming and suppression under chronic low oxygen conditions (50). These effects underscore the brain’s dual role as both a regulator and a target of hypoxic stress.

The fish brain exhibits a high capacity for plasticity, including processes for cellular growth and structural reorganization. Adult neurogenesis, or the creation of new neurons during adulthood, is supported by numerous proliferative stem cell niches distributed along the midline axis (51). These niches contain heterogeneous neural stem/progenitor populations, such as radial glia–like and neuroepithelial-like cells, that sustain ongoing proliferation, differentiation, and neuronal integration across life (52, 53). Emerging evidence further suggests that hypoxia can influence this plasticity by modulating neural cell proliferation. In adult zebrafish, acute hypoxia increases proliferative activity in specific brain regions, including the ventral telencephalon and cerebellum (48), whereas in anoxia-tolerant crucian carp, prolonged anoxia followed by reoxygenation does not alter telencephalic proliferating cell nuclear antigen (PCNA) cell counts, but does induce transient cell death and delayed upregulation of proliferation-related transcripts. This is consistent with a recovery-associated repair response rather than an immediate mitotic increase (54).

In parallel, oxygen limitation induces neurovascular adaptations that contribute to cellular and structural remodeling of the brain, reflected in changes in cerebral blood flow, molecular signaling pathways, and angiogenic responses. Anoxia increases cerebral blood flow in hypoxia-tolerant fish (39, 55, 56), and chronic hypoxia exposure in the brains of goldfish can alter pathways linked to endothelial function, ion channel activity, glutamatergic signaling, and ATP utilization (33, 57), although such responses may vary across fish species. Hypoxia can also activate angiogenic signaling pathways, including vascular endothelial growth factor (VEGF), with evidence of hypoxia-induced neovascularization in adult zebrafish CNS tissue (58). In addition, neural stem cells and endothelial cells share VEGF/VEGF-receptor signaling mechanisms in the adult zebrafish brain, suggesting a potential basis for neurovascular coordination (59). Together, these findings indicate that hypoxia can reshape teleost brain plasticity through effects on proliferation, cell survival, metabolism, and vascular function.

The speckled sanddab (*Citharichthys stigmaeus*), a benthic flatfish common in Elkhorn Slough, California, provides an ecologically relevant model for studying brain responses to hypoxia. Speckled sanddabs utilize the Elkhorn Slough as a nursery habitat during their juvenile stage, before occupying shallow soft bottom habitats along the open coast in the Monterey Bay, CA (6). This species experiences pronounced and recurrent fluctuations in oxygen availability driven by tidal dynamics, local habitat structure, and diel cycles of primary production and respiration of marine macrophytes, which is strongly influenced by eutrophication (10, 60). Field sampling indicates that flatfish presence in the estuary declines strongly with decreasing dissolved oxygen, with fish often being absent below ∼4 mg O₂/L in daytime measurements; notably, nighttime oxygen levels are often lower and anoxia is common in tidally restricted areas (6). Experimental studies further show that juvenile sanddabs activate coping responses under relatively mild reductions in oxygen during acute exposure, including increased ventilation and shifts toward anaerobic metabolism (16), indicating sensitivity to early or moderate oxygen limitation. At the behavioral level, hypoxia impairs critical behaviors such as predator avoidance and escape responses (61), suggesting that these compensatory mechanisms do not fully preserve organismal performance. Despite these ecological and organismal responses, the consequences of hypoxia for the brain and its function remain poorly understood at the cellular (neuroplasticity), neurovascular (morphological remodeling), and molecular levels.

Here, we investigate the effects of prolonged oxygen deprivation (7 days) on the brain of *C. stigmaeus* by integrating histological analyses of neural proliferation, neurovascular remodeling, and molecular responses at the transcriptomic and proteomic levels. For the assessment of neural proliferation and neurovascular changes, we focused on two well-established neurogenic brain regions: the hypothalamic nucleus recessus lateralis (NRL) and the optic tectum. In teleosts, these regions are associated with metabolic regulation, sensory processing, and widespread adult neurogenesis supported by conserved proliferative niches along the midline axis (51, 62, 63). The optic tectum represents a conserved germinal niche contributing to lifelong brain growth across teleost species (63, 64). This integrative approach enables us to examine neural, vascular, and molecular processes in the brain under prolonged hypoxia, addressing a key gap in understanding how environmentally relevant oxygen variability shapes brain plasticity and to identify potential mechanisms of neural adaptation in an ecologically relevant non-model species.

## Materials and Methods

### 4.1 Fish collection and husbandry

Speckled sanddabs occur from Alaska to Baja California, primarily in sandy and soft-bottom habitats in coastal areas and bays (65). Speckled sanddabs settle nearshore in spring and summer, often using bays and estuaries as a nursery habitat for the first year of life. For this project, juvenile speckled sanddabs (*Citharichthys stigmaeus*) were collected from Elkhorn Slough, California, as previously described (16). Fish were captured by beach seine or otter trawl and transferred to aerated coolers with battery-powered airstones for transport. All fishes were then housed for 12 months at Marine Moss Landing Laboratories (MLML), where they were held under a natural light : dark cycle and ambient flow-through seawater conditions prior to the commencement of experiments. Fish were fed daily to satiation using a mix of chopped squid, shrimp, and worms. During experimental periods, fish were fasted 24–36 hours prior to manipulation or tissue collection. All experiments were performed in accordance with institutional guidelines approved by the San Jose State University Institutional Animal Care and Use Committee (*Protocol #1081*).

### 4.2 Experimental hypoxia exposure

Fish were exposed to controlled dissolved oxygen conditions for 7 days, across two experiments. In the first experiment, designed to assess neural cell proliferation, survival, and vasculature, animals were assigned to one of three treatment groups (n = 7 per group): normoxia (8 mg O₂/L), low oxygen (4 mg O₂/L), or hypoxia (2 mg O₂/L). Analyses of neural cell proliferation and survival included all three treatments, whereas vasculature analyses were designed to focus on the contrast between normoxic and hypoxic conditions (8 vs 2 mg O₂/L). In marine systems, environmental hypoxia is typically defined as oxygen concentrations below ∼2 mg O₂/L, whereas higher levels may still induce sublethal physiological stress (<4.5 mg O₂/L in fish; (3)). In the second experiment, designed for transcriptomic and proteomic analyses, animals (n = 7 per group) were exposed to either normoxia (8 mg O₂/L) or hypoxia (2 mg O₂/L) for 7 days. Fish (mean mass = 49 g ± 14.2 SD; age 18–24 months) were randomly assigned to treatment conditions for each experiment.

Hypoxic conditions were created by manipulating the concentration of dissolved oxygen (DO) to desired set points, as previously described (11). Briefly, offshore seawater was pumped through a series of sand filters and settling tanks into a 2,000 L reservoir tank, temperature was stabilized at 12°C using aquarium chillers, and oxygen levels were maintained near 100% saturation (∼8.0 mg O_2_/L) via aeration. The reservoir water was then distributed to the different DO treatment tanks, each holding 500 L of seawater. Bubbling of nitrogen gas (N_2_) was then used to extract oxygen from the water to achieve the desired dissolved oxygen concentrations. Gas flow was managed through solenoid valves controlled by WitroxView software, while oxygen levels were continuously tracked using optical dissolved oxygen probes from Loligo Systems. The conditioned water was then directed into 80 L experimental tanks at a flow rate of 20 ml/s using a single-pass system. Subsequently, sanddabs were placed randomly in 110-gallon tanks (n = 7 per tank) according to the specific treatment groups. Each tank was covered with plastic mesh to prevent escape.

### 4.3 BrdU and EdU administration

To label dividing brain cells, we used 5-bromo-2′-deoxyuridine (BrdU) and 5-ethynyl-2-deoxyuridine (EdU), synthetic thymidine analogs that incorporate into newly synthesized DNA during the S-phase of the cell cycle. Both BrdU and EdU were resuspended in sterile 0.9% NaCl and injected intraperitoneally at 10 mg/kg. For brain cell survival assays, BrdU was administered daily for three consecutive days before the start of hypoxic stress exposure, while EdU was administered 24 and 12 hours before sacrifice to label recently proliferated cells.

### 4.4 Tissue processing and immunohistochemistry

Neural cell proliferation, survival, and vasculature structure in the brain in response to hypoxia were analyzed by immunohistochemistry. Brain tissues were processed for immunofluorescent detection of thymidine analog incorporation and vascular markers. Fish were euthanized using (500 mg/L) of MS-222 (tricaine methanesulfonate) and transcardially perfused with 5 mL of 1M Phosphate-Buffered Saline (PBS). Brains were dissected and stored in 4% paraformaldehyde at 4°C for 48 hours, then dehydrated by immersion in 30% sucrose solution for at least an additional 48 hours. Subsequently, serial coronal sections of the entire brain (40 *µm*) were collected using a freezing sliding microtome (Leica SM2010R) and stored in cryoprotective medium (40% PBS, 30% glycerol, 30% ethylene glycol) at -20 °C.

Immunohistochemistry was performed on free-floating sections using standard protocols (66). For BrdU labeling, all sections were pre-treated with 3 M hydrochloric acid for 15 minutes at 37 °C before incubation with primary antibody. Immunostaining primary antibodies included: rat anti-BrdU (1:2500, Abcam, Cat# ab6326), and lectin tomato (1:200, Vector Laboratories, Cat# DL11741), and Glucose Transporter Type 1 (GLUT-1) to assess vascular morphology (1:200, Novus Biologicals, Cat# NB300-666). EdU detection was performed using the Click-iT™ Plus EdU Alexa Fluor® 555 Imaging Kit (ThermoFisher, Cat# C10638). Fluorescent secondary antibodies (Alexa Fluor® 488, 555, or 647; ThermoFisher) were raised in donkey and used at 1:200 dilution for 2 hours at room temperature in TBS-T (0.01 M Tris-HCl pH 7.4, 0.15 M NaCl, 0.05% Tween 20). Nuclei were counterstained with DAPI (1:2000, Sigma, Cat# D9542). Brain slices were mounted on microscope slides and coverslipped using Molecular Probes™ ProLong™ Diamond Antifade Mountant (Fisher, Cat# P36961).

### 4.5 Imaging and quantitative analysis

To assess brain cell proliferation, survival, and brain vasculature structure in response to different concentrations of DO, immunofluorescent brain sections were imaged and analyzed by observers blinded to the treatment groups. Micrographs were acquired using an Axio Observer 7 fluorescence microscope (Carl Zeiss Microscopy, LLC, USA) and Zen Blue software. All images used for quantification were captured at 20× magnification. Exposure times were optimized for each antigen to avoid signal saturation and were kept constant across all groups. Z-stack images were processed and analyzed using the National Institutes of Health (NIH) ImageJ2 software (v. 2.14.0). Sections were evaluated for tissue integrity and successful EdU/BrdU incorporation. Samples not meeting predefined quality criteria, such as tissue damage (i.e. tears or folds) or poor staining/labeling, were excluded from specific analyses prior to statistical testing.

#### Cell proliferation and survival

Quantification of cell proliferation and survival in the hypothalamic nucleus recessus lateralis (NRL) and tectum of the speckled sanddab was performed manually. Regions of interest were identified based on visible anatomical landmarks and alignment with neurogenic zones described in other teleost species, including zebrafish, *Astatotilapia burtoni*, and *Apteronotus leptorhynchus* (67–70). Cell counts were corrected for the number of slices in the hypothalamus or for measured area in the tectum. For each structure, 4–6 representative coronal slices were analyzed, with two to four images acquired per section.

#### Vascular analysis

Vascular analysis was performed to compare structural vascular features between normoxic (8 mg O₂/L) and hypoxic (2 mg O₂/L) conditions. Quantification of vasculature was performed in two hypothalamic regions per animal: the dorsolateral nucleus (NDL) and the posterior periventricular nucleus (NPP). For each region, three to four coronal sections per animal were analyzed, with non-overlapping fields of view captured per section, yielding six images per region per animal. For vessel diameter and tortuosity, two measurements were obtained per image. Diameter was defined as the perpendicular distance between vessel edges, and tortuosity was calculated as the ratio of vessel segment length to straight-line endpoint distance, following (71). Vessel area was quantified as the percentage of GLUT-1–positive area per image, as GLUT-1 labels endothelial cells of the blood–brain barrier and enables assessment of vascular morphology.

### 4.6 RNA extraction and sequencing

To characterize hypoxia-induced changes in gene expression in the speckled sanddab brain, whole hemibrain tissue was processed for RNA extraction and high-throughput sequencing. Fish were euthanized using (500 mg/L) of MS-222 (tricaine methanesulfonate) and transcardially perfused with 5 mL of 1M Phosphate-Buffered Saline (PBS). Brains were dissected and each half was stored at -80 °C for further processing.

#### RNA Extraction

Total RNA was extracted from whole hemibrain tissue using RNeasy Midi kits (Qiagen, Cat# 75144) according to manufacturer’s instruction and quantity was determined using NanoDrop 2000 (Thermo Fisher Scientific, Waltham, MA, USA). RNA was then submitted and further assessed for quality by Novogene Co. Inc. (Sacramento, US; https://en.novogene.com/) prior to library preparation and sequencing.

#### Library preparation and sequencing

Total RNA quality was assessed prior to library preparation, and only samples with RNA Integrity Number (RIN) > 8 were used. RNA libraries were prepared from 1 μg of total RNA per sample using the NEBNext® Ultra™ RNA Library Prep Kit for Illumina® (NEB, USA).

Library preparation, clustering, and sequencing were performed by Novogene Corporation Inc. (Sacramento, CA, USA) following standard manufacturer protocols. Libraries were sequenced on an Illumina NovaSeq 6000 platform (Illumina, San Diego, CA, USA) using a paired-end 150 bp (PE150) sequencing strategy. Raw reads were demultiplexed and initially quality-filtered by Novogene using fastp. Additional adapter removal and quality trimming were performed using Trimmomatic v0.39 as described below. Read quality was assessed using FastQC (v0.20.0).

### 4.7 De novo transcriptome assembly and differential expression analysis

Because the whole genome sequence for the speckled sanddab is not yet available, a *de novo* transcriptome assembly was generated from quality-trimmed paired-end (PE) reads using the De novo RNA-Seq Assembly Pipeline (DRAP) v1.92, with Oases as the assembler (72). Briefly, raw reads were quality trimmed with Trimmomatic v0.39 (73) run in paired end mode with parameters phred33, ILLUMINACLIP:TruSeq-3-PE.fa:2:30:10:2:TRUE, SLIDINGWINDOW:4:5, LEADING:5, TRAILING:5, MINLEN:25. These trimmed reads were then used for the runDRAP program, skipping the inbuilt trimming step. The resulting transcriptome had an N50 of 2,437 and contained 222,985 contigs. Quality was assessed using BUSCO (Benchmarking Universal Single-Copy Orthologs) with the Actinopterygii odb10 database (74), showing 88.4% completeness (3219/3640), with 44.2% of orthologs (1609/3640) represented only once. The transcriptome was annotated via sequence homology using DIAMOND v2.0.8 against the UniProtKB/Swiss-Prot database (75) (The UniProt Consortium 2023). Annotation was performed twice. Once with only the top match reported (Annotation 1), and again with all matches reported. This second annotation run was filtered in R to select only contigs that aligned to at least one gene from a species within Actinopterygii, with the top match of those alignments assigned as the annotation (Annotation 2). Quality-trimmed reads were aligned to the transcriptome using Salmon v1.2.1 (76) to obtain transcript-level abundances. Transcript-to-gene mapping was then performed using tximport, which aggregated counts from contigs annotated as the same gene (77), resulting in a gene-level count matrix of 19,091 annotated genes for Annotation 1 and 3,319 for Annotation 2. Differential expression analysis was performed on aggregated Annotation 2 counts using the limma package (78) after first filtering low expression genes with the filterByExpr function of edgeR (79) and then applying the voom transformation (80) to model the mean–variance relationship in the log_2_-CPM transformed data. A linear model was fit for each gene, and significance was assessed using empirical Bayes moderation. Genes with P-values < 0.1 and absolute log_2_ fold change ≥ 0.1 were considered significantly differentially expressed.

### 4.8 Proteomic analysis

To characterize protein-level responses to hypoxia, quantitative mass spectrometry-based proteomic analysis was performed on whole hemibrain samples. Sample preparation, LC-MS/MS acquisition, and primary data processing were conducted at the Stanford University Mass Spectrometry Facility (Vincent Coates Foundation Mass Spectrometry Laboratory, Stanford University). Twelve speckled sanddab brain tissue samples were lysed with 500 μL of 10% sodium dodecyl sulfate (SDS)/100 mM triethyl ammonium bicarbonate (TEAB) added for protein extraction and 1-5 μL of Pierce universal nuclease added for DNA disruption. Ceramic beads were added as needed, and the samples were shaken on a bead beater in 5-10 second cycles, until homogenized, and centrifuged at 10,000 RPM for ten minutes at 4 °C. A 5 μL aliquot of supernatant from each sample was transferred into new 1.7 mL Eppendorf tubes, then diluted to a final volume of 50 μL using 10% SDS/100 mM TEAB buffer. The samples were reduced with 10 mM dithiothreitol (DTT) for 20 minutes at 55 °C, cooled to room temperature, and alkylated with 20 mM iodoacetamide for 30 minutes at room temperature in the dark. The samples were acidified to a pH of approximately 1 by adding 2.6 μL of 27% phosphoric acid, followed by the addition of 350 μL of suspension trap (S-trap) loading buffer (90% methanol with 10% 1 M TEAB). The samples were then loaded onto a S-trap plate (Protifi, USA) and washed sequentially with 200 μL aliquots of successive wash solutions (90% methanol with 10% 100 mM TEAB, 90% methanol with 10% 20 mM TEAB, and 90% methanol with 10% 5 mM TEAB). Digestion was performed at 47 °C for two hours using 600 ng of mass spectrometry-grade Trypsin/LysC mix (Promega, USA). The digested peptides were eluted with two 50 μL aliquots of 0.2% formic acid in water, followed by two additional 50 μL aliquots of 50% acetonitrile with 0.2% formic acid in water, into a 96-well collection plate. Finally, the samples were dried using a SpeedVac (ThermoFisher Scientific, USA) and reconstituted with 0.015% n-Dodecyl β-D-maltoside (DDM; Sigma USA) in 0.1% formic acid to a concentration of 50 ng/μL, as determined by the Pierce fluorometric peptide quantitative assay (ThermoFisher Scientific, USA).

#### LC-MS/MS Analysis

Samples were analyzed on the timsTOF Ultra (Bruker Daltonics, Germany) coupled to a nanoElute 2 (Bruker Daltonics, Germany) with 25 ng sample input. Samples were eluted off a PepSep Ultra C18 column (25 cm length x 75 μm ID x 1.5 μm particle size, P/N 1893484; Bruker Daltonics, Germany) at 50 °C connected to a fused silica 10 μm emitter (Bruker Daltonics, Germany) inside a nanoelectrospray Captive Spray source (Bruker Daltonics, Germany) with a 60 min active gradient (2-26% Solvent B in 48-mins, 26-35% Solvent B in 12 mins, 35-95% Solvent B in 0.10 mins, wash at 95% for 7.90 mins; Solvent B: 0.1% formic acid in acetonitrile, Solvent A: 0.1% formic acid in water). The TIMS 1/K0 mobility range was 0.75-1.30 V·s/cm 2, with a ramp time of 100 ms and 100% duty cycle. The source was set to 1600 V, 3.0 l/min dry gas, and 200°C dry temp. diaPASEF windows were created with py_DIA (Skowronek et al. 2022) with a mass range of 350-1250 Da and a mobility range of 0.60-1.45 1/K0 giving an estimated cycle time of 2.25 s. The collision energy ramped from 20 to 59 eV at 0.60 to 1.60 V·s/cm 2.

#### Data Analysis

The .d files acquired in data-independent mode were analyzed using Spectronaut v 19.7 (Biognosys AG, Schlieren, Switzerland) with the directDIA+ feature against the Uniprot pleuronectoidei protein database. Proteolysis with Trypsin was assumed to be specific, allowing for N-ragged cleavage with up to two missed cleavage sites, with a PSM false discovery rate (FDR) of 1%. Data filtering was based on the Q Value, and global normalization was utilized. Cysteine modified with iodoacetamide was set as a fixed modification. Variable modifications included oxidation of methionine and tryptophan, deamidation of asparagine and glutamine, and phosphorylation of serine, threonine, and tyrosine, acetylation of N-terminal and lysine, and pyro-glutamic acid conversion from glutamine and glutamic acid.

### 4.9 Functional enrichment analysis

Gene Ontology and Reactome enrichment analyses were conducted using g:Profiler (version e113_eg59_p19_f6a03c19) via the g:GOSt tool (81). Separate analyses were performed for differentially expressed genes (RNA-seq) and differentially abundant proteins (proteomics). Due to the lack of a reference genome for speckled sanddab (*Citharichthys stigmaeus*), gene identifiers were mapped to human orthologs using UniProtKB/Swiss-Prot annotations derived from DIAMOND homology searches, and enrichment was performed using *Homo sapiens* reference background. For the RNA-seq analysis, 178 unique genes with unadjusted *P* < 0.05 (ranked by nominal p-values from limma-voom) were submitted for enrichment. For the proteomic analysis, Spectronaut candidate protein groups (PG.IsCandidate = TRUE) were mapped to gene symbols where available, and 63 unique annotated proteins were used as input for enrichment analysis. All available Gene Ontology (GO:BP) and Reactome annotations were included. Term sizes were restricted to 100–5000 genes. Significantly enriched terms were defined using g:Profiler’s g:SCS correction for multiple testing, with a significance threshold of *P* < 0.05. This threshold was selected to capture broader transcriptomic patterns for pathway-level analysis, as more stringent multiple testing correction at the gene level resulted in a limited gene set and reduced statistical power. This approach is consistent with established practices in pathway enrichment analyses, where error control is applied at the level of functional categories rather than individual genes (82, 83). Furthermore, overly stringent statistical filtering can exclude genes with modest fold changes that nevertheless contribute to coordinated biological processes (84). This consideration is particularly relevant in *de novo* transcriptome analyses, where annotation uncertainty and reduced mapping efficiency can further limit the number of genes passing strict correction thresholds.

### 4.10 Statistical analysis

Cell proliferation and survival data were analyzed using one-way ANOVA with the factor of DO treatment, followed by Fisher’s LSD post-hoc tests, or Kruskal–Wallis tests with Dunn’s post-hoc comparisons when assumptions were violated. Vascular comparisons were analyzed using unpaired two-tailed Student’s *t*-tests. Transcriptomic differential expression was performed using limma-voom and edgeR pipelines in R (v4.3.2).

Downstream analyses and data visualization were conducted using ggplot2, ggrepel, pheatmap, dplyr, scales, and stringr. Proteomic differential abundance was assessed using two-sided *t*-tests in Spectronaut. Spectronaut candidate proteins were defined using the PG.IsCandidate = TRUE field in the protein-level report. For visualization, log₂ fold change was calculated as mean hypoxia log₂ protein abundance minus mean normoxia log₂ protein abundance, and nominal p-values were used to display the volcano plot; Spectronaut candidate status was used as the primary classification for differentially abundant proteins. Additional statistical analyses and figure preparation were performed in Python (v3.10) and GraphPad Prism (v10.2.2). Statistical significance was set at *P* < 0.05 unless otherwise specified.

### 4.11 Data and code availability

RNA-seq data (*de novo* transcriptome and raw reads) will be deposited in GEO (accession pending). Proteomics data deposition will be in ProteomeXchange (accession pending). Code is available upon request.

## Results

### 5.1 Hypoxia increases neural progenitor activation

Neural cell proliferation, survival, and progenitor activation were quantified in the hypothalamic nucleus recessus lateralis (NRL) and optic tectum following 7 days of exposure to normoxia (8 mg O₂/L), low oxygen (4 mg O₂/L), or hypoxia (2 mg O₂/L). EdU⁺, BrdU⁺, and double-labeled EdU⁺/BrdU⁺ cells were assessed by immunofluorescence (Fig. 1A). In the hypothalamic NRL, hypoxia significantly increased EdU⁺ cell density (a marker of cell proliferation) compared to normoxia (148.1 vs. 95.4 cells/mm²; one-way ANOVA, *F*_2,13_ = 5.84, P = 0.0155; Fisher’s LSD, *P* = 0.0138; normoxia n = 6; low oxygen n = 5; hypoxia n = 5), representing a 1.55-fold increase (Fig. 1B). Low oxygen fish did not differ from normoxic fish in EdU⁺ cell density (*P* = 0.6967). BrdU⁺ cell density (reflecting surviving cells) was significantly reduced under hypoxia relative to normoxia (129.3 vs. 381.2 cells/mm²; one-way ANOVA, *F*_2,13_ = 15.74, *P* = 0.0003; Fisher’s LSD, *P* = 0.0007), corresponding to a 2.95-fold decrease. Low oxygen exposure did not significantly affect BrdU⁺ cell density in speckled sanddabs compared to normoxia (*P* = 0.3200). Double-labeled EdU⁺/BrdU⁺ co-localized area, reflecting neural progenitor cell activation, differed significantly among oxygen conditions in the hypothalamus (Kruskal–Wallis, *H*_2_ = 9.96, *P* = 0.007). Dunn’s post hoc testing confirmed a significant increase in neural progenitor cells under hypoxia relative to normoxia (*P* = 0.008), whereas low oxygen did not differ from normoxia (*P* = 0.60). Median co-localized area increased from 15.11% under normoxia to 88.87% under hypoxia (Fig. 1B–C).

**Fig 1.**
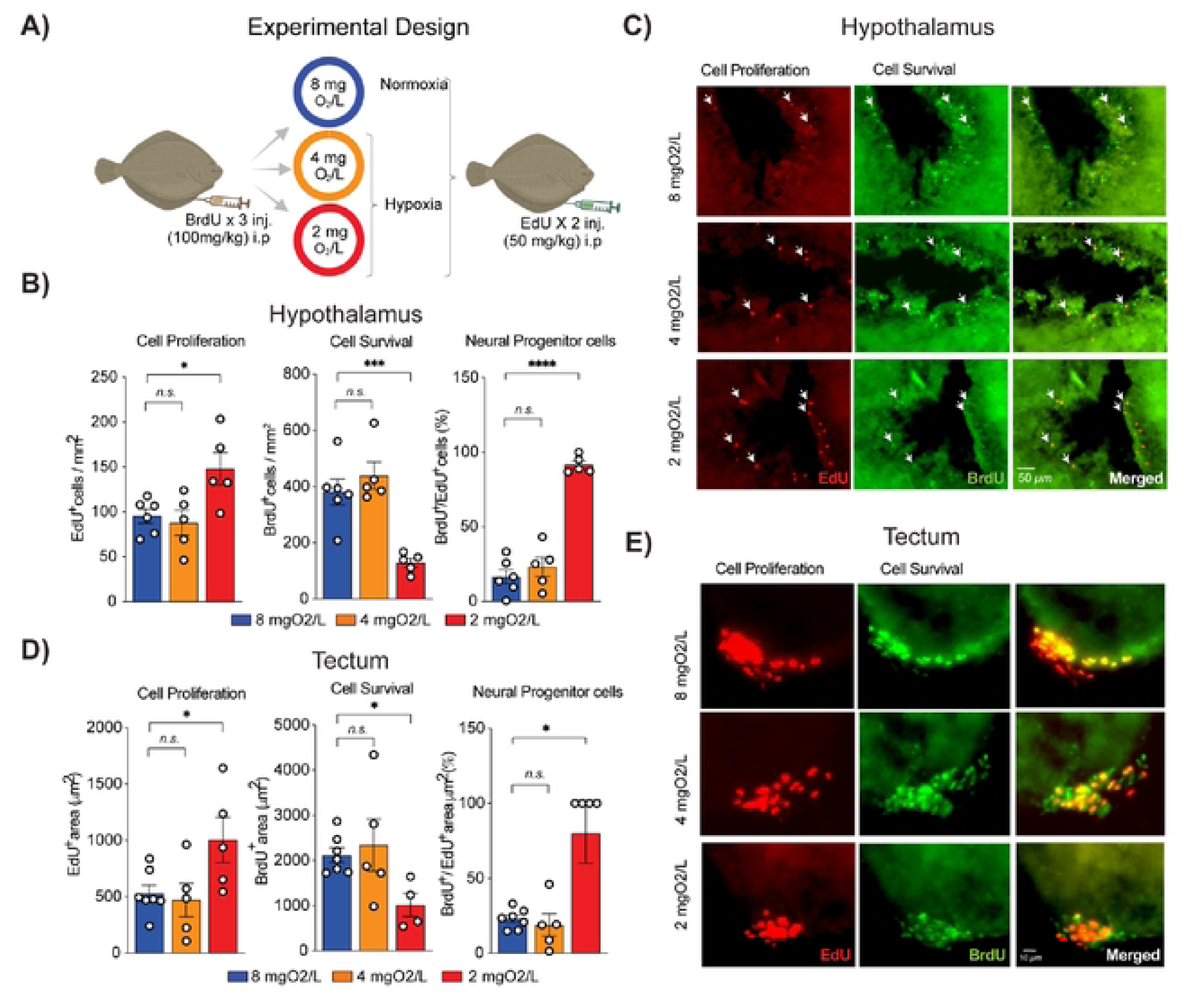
Hypoxia increases cell proliferation and progenitor activation while reducing survival in speckled sanddab brains. (A) Experimental design showing 7-day exposure to normoxia (8 mg O₂/L), low oxygen (4 mg O₂/L), or hypoxia (2 mg O₂/L). BrdU labeled previously divided surviving cells; EdU labeled proliferating cells. (B) Quantification of EdU⁺, BrdU⁺, and EdU⁺/BrdU⁺ cells in the hypothalamic nucleus recessus lateralis (NRL). Bars represent mean ± s.e.m. (n = 5–6 per group). One-way ANOVA with Fisher’s LSD post hoc tests; EdU⁺/BrdU⁺ analyzed by Kruskal–Wallis with Dunn’s post hoc. (C) Representative NRL images showing EdU⁺, BrdU⁺, and double-labeled cells. White arrows indicate colocalized cells. Scale bar = 50 µm. (D) Quantification of EdU⁺, BrdU⁺, and EdU⁺/BrdU⁺ area in the optic tectum (n = 4–7 per group). EdU⁺ and BrdU⁺ analyzed by one-way ANOVA with Fisher’s LSD; EdU⁺/BrdU⁺ analyzed by Kruskal–Wallis with Dunn’s post hoc. (E) Representative optic tectum images showing EdU⁺, BrdU⁺, and colocalized labeling. Scale bar = 10 µm. *P < 0.05; **P < 0.01; ***P < 0.001.

In the optic tectum, hypoxia significantly increased brain cell proliferation in speckled sanddabs. The area of EdU⁺ labeled cells was 1.88-fold higher following exposure to hypoxia compared to normoxia (1003 vs. 533 µm²; one-way ANOVA, *F*_2,14_ = 4.15, *P* = 0.0384; Fisher’s LSD, P = 0.0271; normoxia n = 7; low oxygen n = 5; hypoxia n = 5; Fig. 1D-E). Low oxygen exposed fish did not differ from normoxic fish in cell proliferation (*P* = 0.7522). BrdU⁺ area as a measure of cell survival was reduced under hypoxia relative to normoxia (1019 vs. 2124 µm²; one-way ANOVA, *F*_2,13_ = 3.38, *P* = 0.0659; normoxia n = 7; low oxygen n = 5; hypoxia n = 4; Fig. 1D-E). Although the overall ANOVA was marginally non-significant, pairwise comparison between hypoxia and normoxia reached *P* = 0.0489. Low oxygen did not differ from normoxia in cell survival (*P* = 0.6473). Neural progenitor cells (EdU⁺/BrdU⁺ co-localized area) differed significantly among oxygen conditions (Kruskal–Wallis, *H*(2) = 9.25, *P* = 0.0033; Fig. 1D-E). Dunn’s post hoc test confirmed a significant increase in neural progenitor cells under hypoxia relative to normoxia (*P* = 0.0339), whereas low oxygen did not differ from normoxia (*P* = 0.8412). Median co-localized area increased from 23.4% in normoxia to 100% under hypoxia, corresponding to a 4.27-fold increase (Fig. 1D–E).

### 5.2 Hypoxia induces vascular remodeling

Vascular remodeling in speckled sanddabs was assessed by comparing normoxic (8 mg O₂/L) and hypoxic (2 mg O₂/L) groups following 7 days of exposure (Fig. 2A). In the hypothalamus NRL, hypoxia increased GLUT1⁺ vessel area compared to normoxia (5.46% vs. 4.54%; unpaired two-tailed t-test, *t*_8_ = 2.50, *P* = 0.0369), representing a 1.20-fold increase (Fig. 2B, C). Blood vessel count in the hypothalamus was also elevated under hypoxia (19.38 vs. 15.88; t_8_ = 3.12, P = 0.0142), corresponding to a 1.22-fold increase. In the optic tectum, hypoxia similarly increased blood vessel area (16.36% vs. 12.64%; *t*_8_ = 4.13, *P* = 0.0033) in speckled sanddabs, representing a 1.29-fold increase (Fig 2B, D). However, vessel count did not differ between groups (31.40 vs. 33.07; *t*_8_ = 0.47, *P* = 0.6506) in this region of the brain. No significant differences were observed in vessel diameter, tortuosity, or EdU⁺/GLUT1⁺ co-localization in either region (all *P* > 0.05; Fig. 2B).

**Fig 2.**
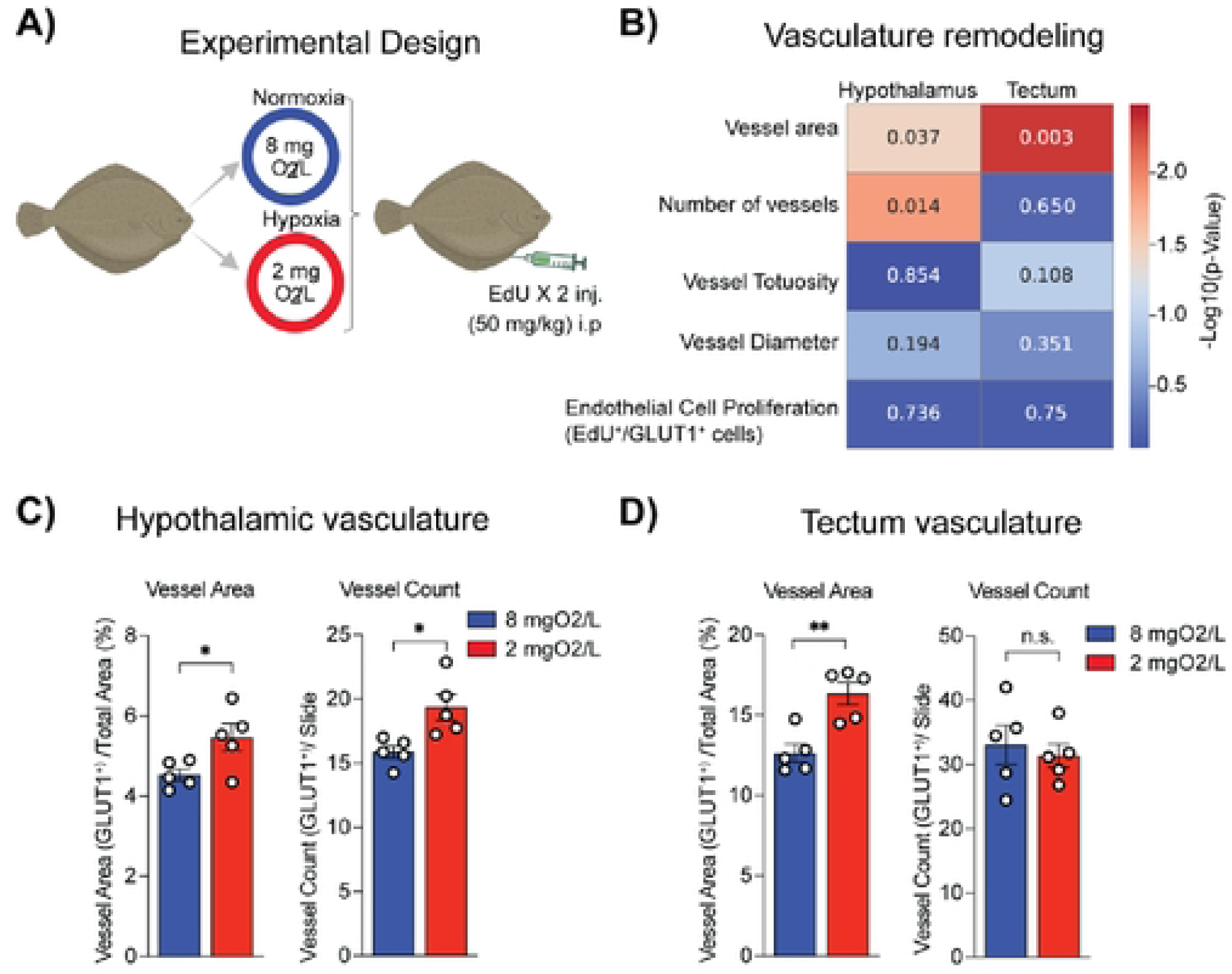
Hypoxia induces vascular remodeling in speckled sanddab brains. (A) Experimental design showing 7-day exposure to normoxia (8 mg O₂/L) or hypoxia (2 mg O₂/L) (n = 5 per group). EdU was administered 24 hours prior to tissue collection. (B) Heatmap summarizing statistical comparisons of vascular remodeling features in the hypothalamus and optic tectum. Values represent two-tailed p-values from unpaired Student’s t-tests. Color scale indicates –log₁₀(p-value). Features analyzed include vessel area, vessel count, vessel tortuosity, vessel diameter, and endothelial cell proliferation (EdU⁺/GLUT1⁺ co-localization). (C–D) Quantification of vessel area (% of total GLUT1⁺ area) and vessel count (GLUT1⁺ vessels per section) in the hypothalamus (C) and optic tectum (D). Data represent mean ± s.e.m. *P < 0.05; **P < 0.01; n.s., not significant.

### 5.3 Hypoxia alters the brain transcriptome

RNA sequencing was performed on whole hemibrains from normoxic (n = 7) and hypoxic (n = 5) speckled sanddabs following 7 days of exposure (Fig. 3A). After removal of low-expression transcripts, 3,253 genes were retained for differential expression analysis. Differential expression identified six genes meeting FDR < 0.1 following Benjamini–Hochberg correction. Using a broader nominal threshold (P < 0.1 and |log₂FC| > 0.1), 552 genes were identified. These genes were used for unsupervised hierarchical clustering, which revealed clear separation between treatment groups (Fig. 3B). For downstream interpretation, a more stringent subset of genes meeting nominal *P* < 0.05 and |log₂FC| ≥ 0.2 was defined. This resulted in 176 differentially expressed genes, including 88 upregulated and 88 downregulated under hypoxia (Fig. 3C).

**Fig 3.**
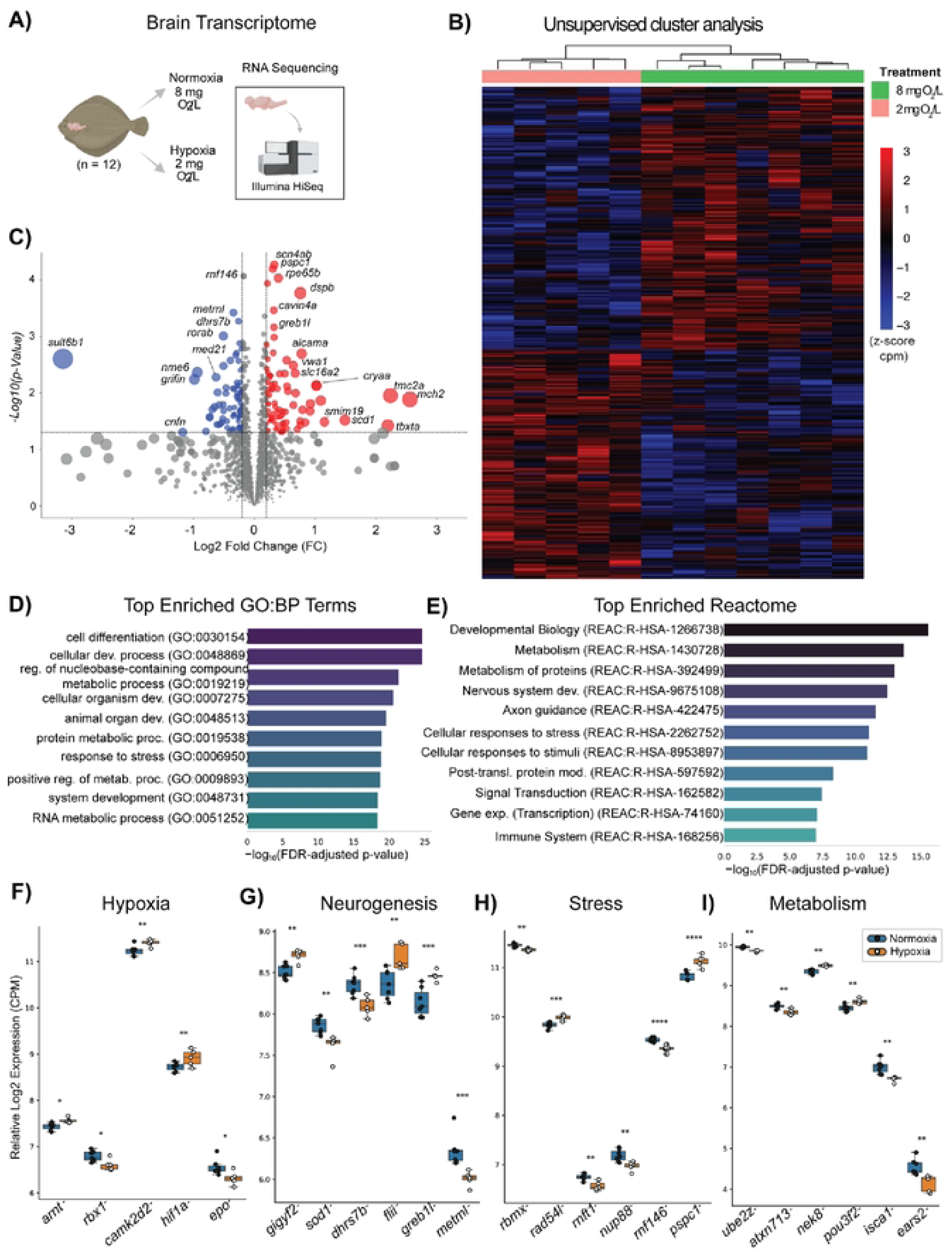
Hypoxia alters the brain transcriptome of speckled sanddabs. (A) Experimental design showing 7-day exposure to normoxia (8 mg O₂/L, n = 7) or hypoxia (2 mg O₂/L, n = 5). Whole hemibrains were collected for RNA sequencing. (B) Unsupervised hierarchical clustering of 552 genes meeting nominal *P* < 0.1 and |log₂FC| > 0.1, showing separation between normoxic and hypoxic transcriptomes. Gene expression values are scaled (z-score) across samples. (C) Volcano plot displaying the same 552 genes. Genes meeting stricter criteria (nominal *P* < 0.05 and |log₂FC| ≥ 0.2; n = 176) are highlighted. (D) Gene Ontology (GO: Biological Process) enrichment analysis of the 176-gene subset. Bars represent –log₁₀(FDR-adjusted *P* values). The top enriched biological processes are shown. (E) Reactome pathway enrichment analysis of the same 176-gene subset. Bars represent –log₁₀(FDR-adjusted P values). (F–I) Expression of selected genes within enriched functional categories. Expression values represent normalized log counts per million (CPM) across individual samples. Panels show (F) hypoxia signaling genes (*camk2d2*, *hif1a*, *epo*, *rbx1*, *arnt*), (G) neurogenesis-related genes (*metrnl*, *dhrs7b*, *greb1l*, *flii*, *sod1*, *gigyf2*), (H) stress-associated genes (*pspc1*, *rnf146*, *rad54l*, *rnft1*, *rbmx*, *nup88*), and (I) metabolism-related genes (*pou3f2*, *isca1*, *nek8*, *atxn7l3*, *ears2*, *ube2z*). Statistical significance was determined using limma-voom differential expression analysis. \**P* < 0.05; **P < 0.01; \*\*\**P* < 0.001; \*\*\*\**P* < 0.0001. Bars represent mean ± s.e.m.

Gene Ontology (GO: Biological Process) enrichment analysis was performed on this 176-gene subset. The most significantly enriched terms included cell differentiation (GO:0030154), cellular developmental process (GO:0048869), regulation of nucleobase-containing compound metabolism (GO:0019219), and multicellular organism development (GO:0007275), among others (adjusted *P* < 1 × 10⁻¹⁹; Fig. 3D; Supplementary Table 1. Reactome pathway enrichment analysis of the same 176-gene subset identified significant enrichment in pathways related to developmental biology (R-HSA-1266738), metabolism (R-HSA-1430728), and protein metabolism (R-HSA-392499), as well as nervous system development and axon guidance (adjusted *P* < 1 × 10⁻¹¹; Fig. 3E; Supplementary Table 1).

To examine transcriptional responses across functional domains, significant genes (*P* < 0.05) were grouped into hypoxia signaling, neurogenesis, stress response, metabolism, and vasculature categories (Fig. 3F–I). Genes were assigned to these categories based on their inclusion in significantly enriched GO: Biological Process and Reactome pathways identified by g:Profiler. From each category, up to six representative genes were selected based on differential expression significance (*P* < 0.05) from the limma-voom analysis. Within the hypoxia signaling panel, *camk2d2*, *hif1a*, and *arnt* were upregulated under hypoxia, whereas *epo* and *rbx1* were downregulated (Fig. 3F). Neurogenesis-associated genes showed mixed expression changes, with *greb1l*, *flii*, and *gigyf2* being upregulated, while *metrnl*, *dhrs7b*, and *sod1* were downregulated (Fig. 3G). Stress-related genes also exhibited both up- and downregulation: *pspc1* and *rad54l* were upregulated, whereas *rnf146*, *rbmx*, *nup88*, and *rnft1* were downregulated (Fig. 3H). Metabolism-related genes displayed predominantly reduced expression under hypoxia, including *isca1*, *atxn7l3*, *ears2*, and *ube2z*, whereas *pou3f2* and *nek8* were upregulated (Fig. 3I).

### 5.4 Hypoxia alters the brain proteome

To investigate protein-level responses to hypoxia, quantitative mass spectrometry–based proteomic analysis was performed on whole hemibrain samples. Spectronaut candidate selection identified 66 candidate differentially abundant protein groups under hypoxia (PG.IsCandidate = TRUE). For downstream annotation and functional enrichment, candidate protein groups were mapped to gene symbols where available, resulting in 63 unique annotated proteins. Protein-level results are reported using gene symbols corresponding to their encoding genes to maintain consistency with transcriptomic and enrichment analyses; however, all values reflect protein-level abundance rather than transcript-level expression. Using a broader nominal threshold (P < 0.02), 105 proteins were selected for unsupervised hierarchical clustering, which revealed separation between normoxic and hypoxic samples (Fig. 4A). To visualize protein abundance changes more broadly, we generated a volcano plot using log₂ fold change and nominal p-values, with Spectronaut candidate proteins highlighted (Fig. 4B). Candidate proteins with higher abundance under hypoxia were shown in red, whereas candidate proteins with lower abundance under hypoxia were shown in blue.

**Fig 4.**
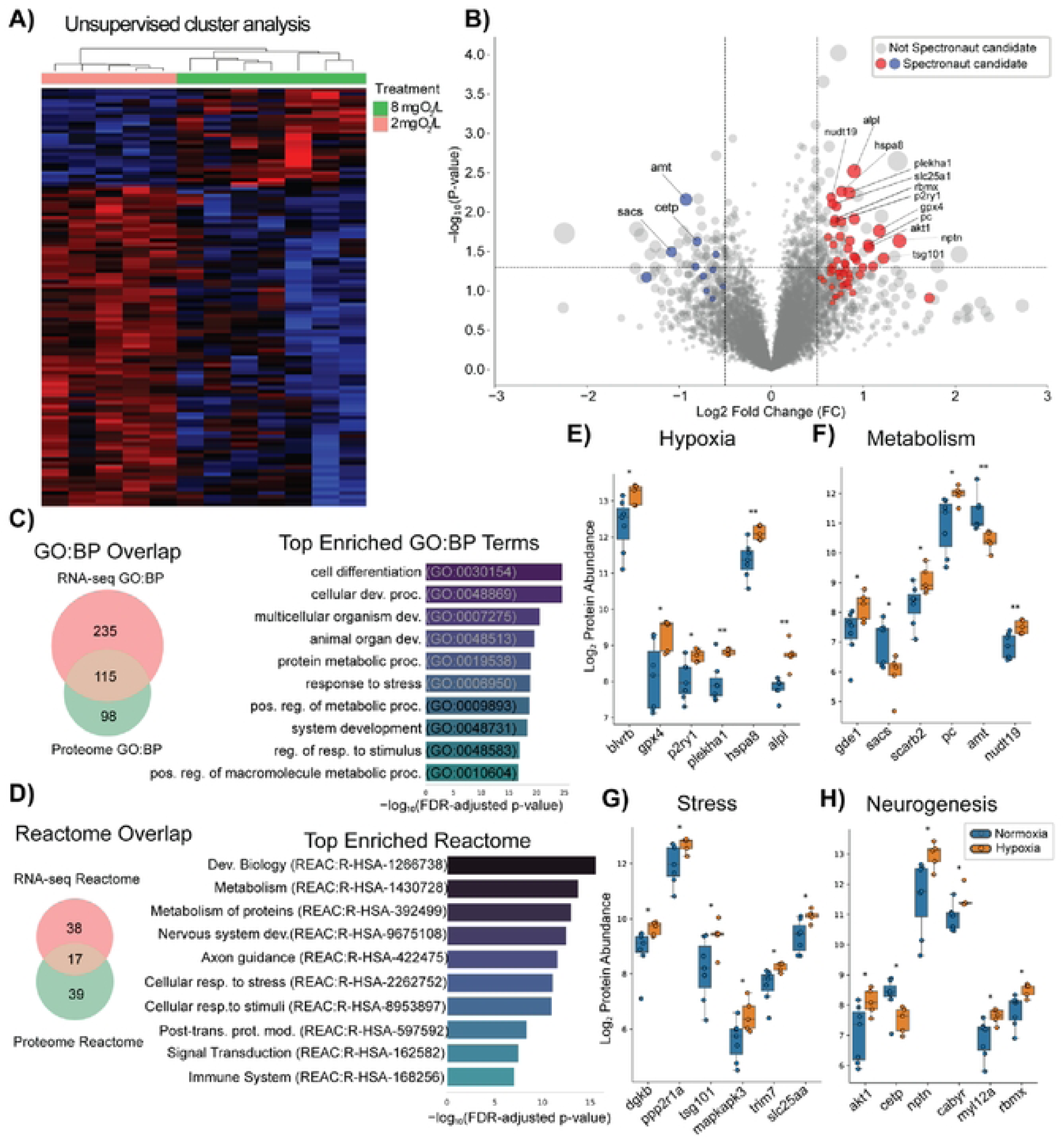
Hypoxia alters the brain proteome and associated functional pathways in speckled sanddab. (A) Unsupervised hierarchical clustering of 105 proteins meeting nominal *P* < 0.02, showing separation between normoxic (8 mg O₂/L) and hypoxic (2 mg O₂/L) samples. Protein abundance values are log₂-transformed and z-score normalized across samples. (B) Volcano plot of protein abundance changes under hypoxia. Each point represents a protein group. The x-axis shows log₂ fold change in protein abundance under hypoxia relative to normoxia, and the y-axis shows −log₁₀ nominal p-value. Proteins identified by Spectronaut as candidate differentially abundant proteins are highlighted, with red indicating higher abundance under hypoxia and blue indicating lower abundance under hypoxia. Non-candidate proteins are shown in gray. Dashed vertical lines indicate |log₂FC| = 0.5, and the dashed horizontal line indicates P = 0.05. Point size reflects combined signal strength based on absolute log₂ fold change and −log₁₀ p-value. (C) Left: Venn diagram showing overlap between GO: Biological Process terms enriched in RNA-seq and proteomic datasets. Right: Bar plot of the top shared GO terms ranked by proteomic enrichment (FDR-adjusted *P* values). (D) Left: Venn diagram showing overlap between Reactome pathways enriched in RNA-seq and proteomic datasets. Right: Bar plot of the top shared Reactome pathways ranked by proteomic enrichment (FDR-adjusted *P* values). (E–H) Protein abundance of selected proteins grouped by functional categories. Expression values represent log₂-transformed protein abundance across individual samples. Panels show (E) hypoxia-response proteins, (F) metabolism-related proteins, (G) stress-associated proteins, and (H) neurogenesis-associated proteins. Proteins were assigned to categories based on enriched GO and Reactome pathways. Statistical significance was assessed using two-sided t-tests. *P < 0.05; **P < 0.01. Error bars represent mean ± s.e.m.

Functional enrichment analysis of the 63 uniquely annotated FDR-significant proteins identified enrichment in Gene Ontology (GO: Biological Process) and Reactome pathways based on human ortholog mapping (Supplementary Table 2). To assess overlap between transcriptomic and proteomic responses, enriched pathways were compared across both datasets. A total of 115 GO: Biological Process terms and 17 Reactome pathways were shared between transcriptomic and proteomic analyses (Fig. 4C–D). Shared GO terms included cell differentiation (GO:0030154), response to stress (GO:0006950), and protein metabolic process (GO:0019538), while shared Reactome pathways included developmental biology (REAC:R-HSA-1266738), cellular responses to stress (REAC:R-HSA-2262752), metabolism of proteins (REAC:R-HSA-392499), axon guidance (REAC:R-HSA-422475), and immune system pathways (REAC:R-HSA-168256).

To examine protein-level changes across functional domains, selected proteins were grouped into hypoxia response, neurogenesis, stress response, metabolism, and vasculature categories (Fig. 4E–H). Proteins were assigned based on their inclusion in enriched GO and Reactome pathways, and up to six representative proteins per category were selected based on nominal P values < 0.05. Within the hypoxia panel, *gpx4*, *p2ry1*, *blvrb*, *plekha1*, *hspa8*, and *alpl* were upregulated under hypoxia (Fig. 4E). Metabolism-associated proteins (*gde1*, *scarb2*, *pc*, *nudt19*) were upregulated, whereas *sacs* and *amt* were downregulated (Fig. 4F). Stress response proteins (*dgkb*, *ppp2r1a*, *tsg101*, *mapkapk3*, *trim7*, *slc25a4*) exhibited upregulated expression under hypoxia (Fig. 4G). Neurogenesis-associated proteins (*akt1*, *nptn*, *cabyr*, *myl12a*, *rbmx*) were upregulated, while *cetp* was downregulated (Fig. 4H).

## Discussion

Overall, chronic hypoxia induced coordinated changes across cellular, vascular, and molecular levels in the brain of *Citharichthys stigmaeus*. At the cellular level, hypoxia increased neural cell proliferation and progenitor activation while reducing survival of newly generated cells. At the tissue level, hypoxia was associated with region-specific vascular remodeling. At the molecular level, transcriptomic and proteomic analyses identified consistent enrichment of pathways related to stress responses, development, and metabolism. Together, these results indicate that hypoxia elicits multi-level responses affecting brain plasticity, vascular structure, and metabolic regulation.

### 6.1 Neural cell proliferation and progenitor cell activation

Seven days of hypoxia increased neural cell proliferation and progenitor cell activation in both the hypothalamic NRL and optic tectum, as indicated by increased EdU labeling and EdU/BrdU co-localization. However, this response was accompanied by reduced survival of newly generated cells, as evidenced by decreased BrdU labeling. These findings indicate that hypoxia stimulates cell proliferation and progenitor activation while reducing long-term survival of newly generated cells. These responses occur within brain regions that play important roles in metabolic regulation and sensory processing and are well-established neurogenic zones in teleosts. In these regions, ongoing progenitor cell activation has been linked to brain growth and lifelong plasticity across species such as the cichlid, *Astatotilapia burtoni*, and ghost knifefish *Apteronotus leptorhynchus* (51, 52, 67–70, 85). Therefore, the present results suggest that hypoxia actively remodels neurogenic niches in the hypothalamus and tectum of speckled sanddabs (*Citharichthys stigmaeus*), potentially reconfiguring circuits involved in energy homeostasis and sensory integration, and thereby contributing to behavioral adaptation to low-oxygen environments (51, 52, 85).

The observed increase in neural cell proliferation is consistent with prior work showing that low-oxygen can regulate neural stem/progenitor cell function, including HIF-1α–linked control of Wnt/β-catenin activity and adult hippocampal neurogenesis processes (86), and that intermittent hypobaric hypoxia can increase hippocampal neuroprogenitor proliferation leading to more newborn neurons in adult rats (87). In teleosts, hypoxia activates neural stem/progenitor and radial glial populations in zebrafish brain and spinal cord embryos, promoting proliferation and subsequent neuronal differentiation (88, 89). In adult zebrafish, acute hypoxia has also been reported to stimulate cell proliferation in telencephalic and cerebellar regions, measured via BrdU-positive cells (48). However, proliferative activation does not necessarily imply long-term persistence of new cells, and the reduced survival of BrdU-labeled cells observed here suggests that newly generated cells may be more vulnerable under hypoxia, potentially reflecting increased apoptotic vulnerability. This is consistent with evidence of increased apoptosis and reduced neuron numbers in salmon spinal cord following early post-hatching hypoxic exposure (90), and with broader vertebrate reviews describing oxygen-level- and duration-dependent “dual effects” of hypoxia on neural stem/precursor populations (91, 92).

These results suggest that hypoxia does not simply enhance cell proliferation, but instead shifts the balance between cell production and survival. In teleosts, adult neurogenesis is inherently characterized by high rates of cell turnover, with many newly generated cells undergoing programmed cell death (70, 93). The pattern observed here, elevated proliferation alongside reduced survival, indicates that hypoxia may amplify this intrinsic turnover process, accelerating cycles of cell production and elimination rather than increasing overall cell addition. Such a mechanism would allow for continued plasticity while preventing excessive or maladaptive cell accumulation under low-oxygen conditions. However, the underlying basis of this response remains unclear, and may reflect selective survival of more hypoxia-tolerant cell populations, altered cellular phenotypes, or increased replacement of vulnerable cells. Consistent with this interpretation, the increase in EdU⁺/BrdU⁺ co-localization observed here, in the absence of increased overall BrdU⁺ cell survival, suggests enhanced progenitor activation without sustained persistence.

Exposure to 4 mg O₂/L did not produce detectable changes in cell proliferation or survival, suggesting that the threshold for cell-level neurogenic responses in *C. stigmaeus* lies below this level after 7 days of exposure. Notably, 2 mg O₂/L corresponds to the commonly used operational definition of environmental hypoxia in coastal marine systems (94), but physiological and behavioral effects in fish are known to occur across a broader and species-dependent range of oxygen conditions (3, 95). Thus, the absence of detectable neurogenic effects at 4 mg O₂/L should not be interpreted as an absence of biological impact more broadly. Previous literature has shown that sublethal effects on fish growth and consumption become consistently negative below approximately 4.5 mg O₂/L and that mean thresholds for reduced growth and food consumption across multiple fish datasets cluster around ∼5–6 mg O₂/L, values substantially higher than the 2–2.5 mg O₂/L definition often used for environmental “hypoxia” in marine systems (96). More broadly, reviews emphasize that the upper DO threshold used to label waters “hypoxic” varies among studies (often cited as ∼2–5 mg O₂/L) and that the 2 mg O₂/L convention can underestimate the onset of hypoxic impacts for many taxa (95).

It remains unclear whether hypoxia preferentially affects neuronal, radial glial (astroglia-like), or other progenitor populations in *C. stigmaeus*, or whether any hypoxia-associated newborn cells survive long enough to integrate into established circuits. In teleosts, progenitor populations in these regions—often including radial glia—can generate newborn cells that differentiate into neurons, with lineage studies also showing contributions to glial fates (e.g., oligodendrocytes and radial glial cells) (62, 97). However, further work is needed to determine how hypoxia shapes these cell populations in *C. stigmaeus,* including whether the observed responses reflect selective generation or survival of specific cell types or broader changes in cell turnover and incorporation in to brain system. Future studies combining lineage tracing with longer survival time points will be essential to resolve these questions.

### 6.2 Neurovascular remodeling under hypoxia

Hypoxia can challenge brain homeostasis by constraining oxygen delivery while neural tissue maintains high metabolic demand. In the present study, seven days of hypoxia (2 mg O₂/L) produced region-specific vascular morphological adaptations in the speckled sanddab brain. This remodeling was characterized by increased vessel area in the hypothalamus and optic tectum and increased vessel number in the hypothalamus, without detectable changes in vessel diameter, tortuosity, or EdU⁺/GLUT1⁺ endothelial proliferation. Together, these results support a model in which 7 days of hypoxia promotes early microvascular expansion in select brain regions.

The vertebrate brain depends on continuous delivery of oxygen and glucose to sustain neuronal activity, and this regulation is implemented by a multi-cellular neurovascular system that coordinates perfusion with local metabolic demand. Beyond flow regulation, cerebrovascular cells also contribute to broader homeostasis by participating in the cerebral barrier function, signaling crosstalk with glia/immune cells, and pathways involved in metabolite and protein waste clearance (98, 99). A well-established mechanism linking low oxygen availability to vascular remodeling is activation of the hypoxia-inducible factor (HIF) pathway, which induces transcriptional programs that promote angiogenesis and metabolic adaptation (100). Previous work shows that hypoxia-driven angiogenesis is strongly shaped by pro-angiogenic growth factor signaling, especially VEGF family pathways, and is also influenced by vessel-stabilizing/destabilizing cues such as angiopoietins (including ANGPT2) acting in concert with VEGF (101).

The pattern observed in speckled sanddabs, greater vessel area without changes in average vessel diameter or tortuosity, suggests an early-stage remodeling process that likely occurs at the microvascular/capillary level. In mammalian models, chronic hypoxia is a potent driver of cerebral microvascular “angioplasticity,” with capillary density increases beginning on week one and continuing over subsequent weeks (102). Longitudinal *in vivo* imaging further demonstrates that, in mouse cortex under chronic hypoxia, new sprouts can emerge after ∼7–14 days and later form new capillary connections, providing a comparative template for interpreting why structural change can precede (or be only weakly coupled to) clear proliferation signatures at a single sampling window (103). In fish, hypoxia also robustly activates hypoxia-inducible factor (HIF) signaling and downstream pathways relevant to vascular and metabolic adaptation, although the regional and cellular specificity of adult fish brain vascular remodeling remains less comprehensively mapped than in the mammalian cortex (104).

The more pronounced remodeling response in the hypothalamus may reflect regional differences in physiological demand and integration. In teleosts, hypothalamic and preoptic areas are major neuroendocrine/metabolic integration hubs, and multiple lines of evidence indicate that fish hypothalamic function is sensitive to environmental hypoxia, including documented hypoxia-induced neurochemical and enzyme pathway disruptions in the hypothalamus of hypoxia-tolerant marine fish (105). While our study does not directly test hypothalamic oxygen sensing mechanisms, the regional selectivity of the vascular phenotype is consistent with the broader concept that neurovascular remodeling is targeted to circuits and regions where maintaining substrate delivery is especially consequential for homeostatic regulation (100).

Despite the increase in vascular area and (in the hypothalamus) vessel number, we did not detect a significant rise in EdU⁺/GLUT1⁺ colabeled endothelial cells, indicating no increase in proliferating endothelial cells. This could reflect the timing of EdU delivery relative to the proliferation window, or it may indicate that early remodeling under these conditions is driven by mechanisms that do not require substantial endothelial cell division. One plausible functional overlay is altered endothelial transporter expression: GLUT1 is a canonical blood–brain barrier glucose transporter highly enriched in the brain microvascular endothelium, and hypoxia/HIF signaling can directly regulate GLUT1 gene expression in multiple cell contexts (106). Importantly for interpretation of GLUT1-based vascular readouts, mammalian studies show that chronic hypoxia can increase both cerebral vascularity and the density of capillary GLUT1 immunostaining, linking hypoxia exposure to increased GLUT1 signaling through structural and/or transporter-expression mechanisms (107). In this framework, increased GLUT1 signal in speckled sanddabs could reflect functional adaptation of existing vessels, structural expansion of microvessels, or a combination—an ambiguity that can be addressed in future work by pairing GLUT1 with additional endothelial markers and/or perfusion measures. An additional non-mutually-exclusive mechanism is intussusceptive angiogenesis, in which existing vessels split via intraluminal pillar formation rather than classic sprout-driven growth. Intussusceptive angiogenesis can increase vascular complexity and surface area with minimal endothelial proliferation and can occur rapidly compared with sprouting angiogenesis, making it consistent with microvascular expansion in the absence of elevated EdU labeling (108). Functionally, this form of vascular remodeling may improve oxygen and nutrient delivery by increasing capillary surface area and optimizing blood flow distribution without the energetic cost of extensive endothelial proliferation. In hypoxic conditions, such adaptations could help maintain cerebral perfusion and metabolic support, thereby preserving neuronal function and enhancing survival in speckled sanddabs.

Finally, the vascular consequences of hypoxia are strongly dependent on dose, duration, and organismal context: moderate hypoxia can support compensatory angiogenesis and remodeling, whereas more severe or prolonged hypoxia can undermine microvascular integrity and contribute to blood brain barrier disruption, neuroinflammation, and tissue injury (109). Within the endpoints assessed here, the speckled sanddab response is most consistent with an adaptive remodeling program rather than overt pathological hyperplasia. Future studies should therefore test mechanistic drivers and thresholds by quantifying region-specific expression of HIF targets associated with angiogenesis, including the potential contribution of intussusceptive angiogenesis, and oxygen delivery (e.g., vegf-family ligands, epo, and angiopoietin/Tie2 axis components), and by evaluating whether longer or repeated hypoxia exposures shift remodeling toward more pronounced structural change or toward barrier dysfunction (100).

### 6.3 Integrated transcriptomic and proteomic responses to hypoxia

Hypoxia induced coordinated molecular changes occurred in the speckled sanddab brain at both transcriptomic and proteomic levels, revealing an integrated regulation of stress responses, metabolism, and plasticity under reduced oxygen availability. Although overlap between individual genes and proteins was limited, both datasets revealed consistent enrichment of biological processes related to cell differentiation, response to stress, protein metabolic process, system development, and regulation of response to stimulus, as well as Reactome pathways such as Developmental Biology, Metabolism, Nervous System Development, Axon Guidance, Cellular Responses to Stress, and Signal Transduction. Similar multi-level molecular responses have been reported in other fish systems, where transcriptomic and proteomic analyses identify changes across these major functional domains (110–112). The limited correspondence between transcriptomic and proteomic datasets at the level of individual targets at a single time-point is commonly reported in multi-omic experiments because protein abundance reflects not only transcription of genes, but also RNA processing, translation efficiency, protein turnover, and time-lagged dynamics between these regulatory layers (113). In contrast, convergence at the level of enriched pathways provides a more biologically meaningful view of the data, as pathway-level aggregation reduces noise and highlights coordinated functional responses (114). Together, these shared processes indicate that hypoxia affects multiple functional domains, including stress responses, development, and metabolism.

#### Stress pathways and hypoxia

Enrichment of biological processes such as response to stress, cellular response to stimulus, regulation of response to stress, intracellular signal transduction, and immune system process indicates that hypoxia activates a broad stress-response and homeostasis regulatory response in the brain of the speckled sanddab. Reactome pathways including cellular responses to stress, cellular responses to stimuli, signal transduction, and immune system further support activation of signaling systems involved in cellular survival and adaptation.

Gene-level results showed upregulation of *hif1a* and *arnt*, consistent with activation of the HIF transcriptional complex, a central regulator of hypoxia responses in vertebrates and fish (104, 115). This aligns with enrichment of stress and signaling pathways identified in both datasets. The concurrent increase in *camk2d2* suggests engagement of Ca²⁺-dependent signaling, which is linked to neuronal plasticity and synaptic regulation (116). And, while the specific role of CAMK2D paralogs in teleost hypoxia responses remains under-characterized, the directionality of *camk2d2* regulation in our results is consistent with hypoxia-linked remodeling of neuronal signaling states (117). Downregulation of *rbx1* and *epo* is consistent with altered proteostasis regulation under hypoxic stress. Given the role of *rbx1* in ubiquitin-mediated proteostasis, reduced expression may alter protein turnover dynamics under hypoxia, while decreased *epo* expression likely reflects tissue-specific regulation rather than absence of hypoxia signaling (115, 118).

At the protein level, increased abundance of GPX4, BLVRB, and HSPA8 is consistent with enhanced oxidative stress protection and protein quality control, matching enrichment of stress-response pathways. Together, these results indicate that hypoxia activates a conserved molecular stress-response program integrating oxygen sensing, redox balance, and cellular maintenance. BLVRB participates in heme-derived biliverdin processing linked to antioxidant bilirubin production, GPX4 is a key enzyme preventing lipid hydroperoxide accumulation, and HSPA8/HSC70 is a central chaperone supporting protein quality control and selective autophagy pathways (119). Changes in P2RY1, PLEKHA1, and ALPL suggest a broader remodeling of signaling (120), membrane-associated pathways (121), and potentially neuronal development (122) . Consistent with the central role of mitochondria as both regulators and targets of hypoxic stress, these adaptations align with mechanistic frameworks of hypoxia tolerance that emphasize enhanced redox regulation and proteostatic capacity under fluctuating oxygen conditions (123). In sum, these molecular patterns are consistent with a coordinated hypoxia response that integrates oxygen sensing, oxidative stress protection, and cellular maintenance in the sanddab brain.

#### Developmental pathways and neurogenesis

Enrichment of developmental processes such as cell differentiation, cellular developmental process, system development, and neurogenesis, together with Reactome pathways including developmental biology, nervous system development, and axon guidance, indicates activation of plasticity-related pathways under hypoxia. Enrichment of developmental and neurogenic pathways (e.g., nervous system development, cellular differentiation, axon guidance) is consistent with the observed increase in proliferative activity at the cellular level and supports activation of plasticity-related programs under hypoxia. These molecular signatures align with the increased neural proliferation and progenitor activation observed in the hypothalamic NRL and optic tectum. At the gene level, differential expression of neurogenesis-associated genes (e.g., *greb1l*, *flii*, *gigyf2*, *metrnl*, *dhrs7b*, *sod1*) suggests selective modulation of developmental signaling networks rather than uniform activation of neurogenesis.

At the protein level, increased abundance of AKT1, NPTN, CABYR, MYL12A, and RBMX further supports activation of pathways involved in neuronal growth, cytoskeletal organization, and cellular differentiation. These results are consistent with the enriched developmental pathways and support a link between molecular signaling and cellular proliferation. Mechanistically, hypoxia has been shown to regulate neurogenesis through HIF-dependent signaling and erythropoietin-mediated pathways (124). In teleosts, where adult neurogenesis is widespread, activation of these pathways provides a substrate for structural remodeling (51). Together, these results indicate that hypoxia engages developmental signaling programs that are associated with increased proliferative activity observed at the cellular level.

#### Metabolic pathways and metabolism

Enrichment of metabolic processes such as protein metabolic process, positive regulation of metabolic process, small molecule metabolic process, transport, and homeostatic process, together with Reactome pathways including metabolism, metabolism of proteins, and transport of small molecules, indicates broad reorganization of cellular metabolism under hypoxia. Comparable responses have been reported in fish brain systems, where hypoxia induces pathway-specific metabolic adjustments rather than a global shutdown of energy metabolism (111, 112, 125). For example, transcriptome studies in fish brain under acute or chronic hypoxia repeatedly identify metabolic pathway remodeling as a central response feature (126). At the transcript level, downregulation of genes such as *isca1* (iron–sulfur cluster assembly) and *ears2* (mitochondrial tRNA synthetase), both associated with mitochondrial function, together with changes in regulatory genes including *ube2z* (ubiquitin-conjugating enzyme) and *atxn7l3* (transcriptional co-regulator), suggests modulation of energy production and protein processing pathways. Concurrent upregulation of *pou3f2* (neural transcription factor) and *nek8* (cell cycle and signaling kinase) further indicates activation of regulatory programs that may support cellular adaptation to low oxygen conditions. Comparable brain transcriptomic analyses in fish explicitly report hypoxia-associated mitochondrial dysfunction signatures and modulation of broader metabolic pathways (127). At the protein level, increased abundance of GDE1 (glycerophosphodiester metabolism), SCARB2 (lysosomal membrane protein), NUDT19 (peroxisomal CoA metabolism), and PC (pyruvate carboxylase), together with decreased abundance of SACS (chaperone-like protein) and AMT (glycine cleavage enzyme), indicates selective adjustment of metabolic pathways rather than global suppression. These findings are consistent with multi-omics fish brain hypoxia studies showing pathway-level remodeling across mRNA/miRNA/protein layers and with mechanistic reviews emphasizing that hypoxia adaptation spans coordinated changes in protein synthesis, mitochondrial respiration, and nutrient metabolism (35). Taken together, the combined transcriptomic and proteomic responses support a model of metabolic flexibility, where transcriptional and protein-level regulation operate at different layers to maintain cellular homeostasis under hypoxic stress (111, 115, 123, 125).

## Conclusions

This study provides the first characterization of hypoxia-associated brain plasticity in speckled sanddab, an ecologically relevant but non-model estuarine flatfish. By integrating neural cell proliferation and survival, vascular morphology, transcriptomics, and proteomics, we show that prolonged hypoxia affects the brain across multiple levels of organization. At the cellular level, hypoxia increased proliferative activity while reducing survival of newly generated cells, indicating a dynamic balance between cell production and turnover. At the tissue level, hypoxia promoted region-specific vascular remodeling, suggesting changes in oxygen delivery or vascular organization in metabolically active brain regions. At the molecular level, integrated transcriptomic and proteomic analyses revealed consistent activation of stress-related, developmental, and metabolic processes, despite limited overlap between individual genes and proteins. Together, these findings support a model in which hypoxia drives multi-level reorganization of brain structure and function, linking cellular and vascular brain plasticity with molecular reprogramming. These responses include features consistent with conserved vertebrate hypoxia biology, such as stress signaling, metabolic remodeling, and vascular plasticity, while also demonstrating that hypoxia can alter cellular dynamics in brain regions associated with adult neurogenesis. This work expands current understanding of fish brain responses to hypoxia beyond traditional model species and suggests that brain plasticity may be an important component of responses to fluctuating oxygen conditions in estuarine environments. Although these responses may help maintain brain function under reduced oxygen availability, they may also carry costs, including reduced survival of newly generated cells and potential consequences for neural function. In ecologically relevant contexts such as Elkhorn Slough, increasing hypoxia may therefore affect key behaviors in juvenile flatfish, with consequences for survival, recruitment, and the nursery function of estuarine habitats. These effects could ultimately scale to influence population dynamics and coastal fisheries, underscoring the importance of managing eutrophication and other drivers of hypoxia in these systems.

